# Vinorelbine causes a neuropathic pain-like state in mice via STING and MNK1 signaling associated with type I interferon induction

**DOI:** 10.1101/2023.06.03.543579

**Authors:** Úrzula Franco-Enzástiga, Keerthana Natarajan, Eric T. David, Krish J. Patel, Abhira Ravirala, Theodore J. Price

## Abstract

Type I interferons (IFNs) increase the excitability of dorsal root ganglion (DRG) neurons via activation of MNK-eIF4E translation signaling to promote pain sensitization in mice. Activation of STING signaling is a key component of type I IFN induction. Manipulation of STING signaling is an active area of investigation in cancer and other therapeutic areas. Vinorelbine is a chemotherapeutic that activates STING and has been shown to cause pain and neuropathy in oncology clinical trials in patients. There are conflicting reports on whether STING signaling promotes or inhibits pain in mice. We hypothesized that vinorelbine would cause a neuropathic pain-like state in mice via STING and signaling pathways in DRG neurons associated with type I IFN induction. Vinorelbine (10 mg/kg, i.v.) induced tactile allodynia and grimacing in WT male and female mice and increased p-IRF3 and type I IFN protein in peripheral nerves. In support of our hypothesis, vinorelbine-mediated pain was absent in male and female Sting*^Gt/Gt^* mice. Vinorelbine also failed to induce IRF3 and type I IFN signaling in these mice. Since type I IFNs engage translational control via MNK1-eIF4E in DRG nociceptors, we assessed vinorelbine-mediated p-eIF4E changes. Vinorelbine increased p-eIF4E in DRG in WT animals but not in Sting*^Gt/Gt^*or *Mknk1^-/-^* (MNK1 KO) mice. Consistent with these biochemical findings, vinorelbine had an attenuated pro-nociceptive effect in male and female MNK1 KO mice. Our findings support the conclusion that activation of STING signaling in the peripheral nervous system causes a neuropathic pain-like state that is mediated by type I IFN signaling to DRG nociceptors.

## Introduction

Vinorelbine is a chemotherapeutic agent that belongs to the semisynthetic vinca alkaloid family (1). This drug is FDA-approved for non-small cell lung cancer (NSCLC) treatment (2). Vinorelbine is also used to treat metastatic breast carcinoma, and has shown effectiveness in advanced forms of melanoma and renal cell carcinoma (1). NSCLC and breast cancer are estimated to be the top two cancer incidences in 2023 in the US (3). Despite serving as a first-line treatment in advanced NSCLC (4), vinorelbine has been reported to cause peripheral neuropathy (5-10) and induce pain in cancer patients, mostly at the tumor site or in areas innervated by nerves compressed by the tumor (1).

Vinorelbine mainly acts as a destabilizing agent of microtubules (4, 11). Microtubule destabilizers have been shown to activate stimulator of interferon genes (STING)- interferon regulatory factor 3 (IRF3) dependent production of IFN-β and downstream interferon-stimulated genes (ISGs) in both immune and triple negative breast cancer cells (12). Type I IFNs are powerful monomeric cytokines that induce an antiviral response upon infection through the upregulation of molecules that antagonize viral replication (13). Regulation of type I IFNs by STING depends on the IRF family members IRF3/7 (14). The expression of IRF7 in most cells is low, while IRF3 is ubiquitously expressed triggering the phosphorylation of IRF3 upon viral infection in nearly all cell types (15). Loss-of-function mutation in the IRF3 gene impairs interferon expression in patients, suggesting a critical role of this transcription factor in the antiviral interferon responses (16, 17).

Previously we showed that IFN-α and IFN-β activate interferon receptors (IFNR) localized on DRG nociceptors driving hyperexcitability and nociception in mice (18, 19). Type I IFN-induced nociception in mice depends on mitogen activated protein kinase interacting kinase (MNK1) signaling to elongation initiation factor 4E (eIF4e). Additional evidence from preclinical models (20) and clinical studies (21) supports the pronociceptive actions of type I IFNs. On the other hand, the role of STING in nociception is far from clear. Some findings have shown that STING-mediated induction of type I IFN signaling attenuates nociception via an action in the spinal dorsal horn (22, 23). Other studies show a clear peripheral sensitization effect of STING signaling activation or the attenuation of nerve damage-induced pain using STING inhibitors in rodents (24, 25). Given the clinical use of vinorelbine and reports of neuropathic pain without any mechanistic insight, we sought to test the hypothesis that this chemotherapeutic causes pain via STING and type I IFN signaling. Our work provides clear evidence in support of this mechanism of action of vinorelbine in male and female mice.

## Methods

### Animals

All experiments were carried out in male and female mice. The following mouse strains were used: ICR outbred mice purchased from Envigo Laboratories (4-6 weeks, weighing from 20 to 25 g, RRID: IMSR_CRL:022); *Mknk1^-/-^* (MNK1 KO), a gift from the Sonenberg laboratory at McGill University and their C57BL/6J WT (WT) littermates (26, 27) bred at the University of Texas at Dallas (UTD); Sting*^Gt/Gt^*also known as Goldenticket mice (*Tmem173^Gt^*) and the suggested controls C57BL/6J WT (Lot: 000664) purchased from Jackson laboratory.

Purchased mice were used in experiments starting one week after arrival at the animal facilities at the University of Texas at Dallas. Animals had free access to food and water before the experiments. Mice were housed in groups of 4 per cage in non-environmentally enriched cages with food and water *ad libitum* on a 12 h non-inverted light/dark cycle. All animal procedures were approved by The University of Texas at Dallas IACUC. Experiments were in compliance with the National Institutes of Health Guide for Care and Use of Laboratory Animals (Publication No. 85-23, revised 1985). Animals were monitored for health according to IACUC guidelines at The University of Texas at Dallas prior and during experimentation. This study was not pre-registered.

### Drugs

Vinorelbine (Methyl (2β,3β,4β,5α,12β,19α)-4-acetoxy-15-[(12S,14R)-16-ethyl-12- (methoxycarbonyl)-1,10-diazatetracyclo[12.3.1.03,11.04,9]octadeca-3(11),4,6,8,15- pentaen-12-yl]-3-hydroxy-16-methoxy-1-methyl-6,7-didehydroaspidospermidine-3- carboxylate), a potent STING activator (12, 28), was purchased from TargetMol (Cat. No. T0190). The drug was dissolved in 3% dimethyl sulfoxide (DMSO) in saline.

### Drug administration scheme

Chemotherapeutic drugs are administered to patients using restricted doses. In this work, we administered the maximum tolerated dose (MTD) of vinorelbine (10 mg/kg, *i.v*; (29)) to male and female mice. MTD refers to the highest dose of a drug that causes a therapeutic effect but not unacceptable side effects. The dose 10 mg/kg of vinorelbine was previously reported to be well tolerated after a single-dose or 3 repeated cycles at a single-dose (29). Additionally, the same dose or similar doses have been used in mice in other reports (30-32).

Our drug administration scheme was based on pilot experimental trials, which showed that vinorelbine (10 mg/kg, i.v.) but not vehicle (DMSO 3% in PBS) causes tactile allodynia and facial grimace responses. These effects were aggravated by the second administration on day 7. Based on these pilot results and the previous clinical and preclinical literature, we decided to do two i.v. administrations of vinorelbine separated by a week for the rest of the experiments.

### Western blot

Mice were anesthetized with 2.5% isoflurane and decapitated. Sciatic nerves and lumbar DRGs were dissected and frozen on dry ice. The tissues were stored in −80°C until they were processed for Western blot. Tissues were homogenized in ice-cold lysis buffer (50mM Tris pH 7.4, 150mM NaCl, and 1mM EDTA pH 8.0, 1% Triton X-100; 400 µl for DRGs and 200 µl for sciatic nerve) with phosphatase inhibitor cocktail added to the lysis buffer immediately prior to use. Homogenates were centrifuged (Eppendorf) at 10,621 g for 10 min at 4°C. The extracted protein was collected and stored at −80°C. Protein concentration was quantified by the Bicinchoninic acid (BCA) assay (Cat. #23225, ThermoFisher Scientific). Fifteen micrograms of total protein were resolved by denaturing in 10% and 15% sodium dodecyl sulfate-polyacrylamide gel electrophoresis and were then transferred overnight to polyvinylidene difluoride (PVDF) membranes (Cat. # IPFL00010, Millipore Sigma). 5% bovine serum albumin (BSA) in 1X TTBS (150 mM NaCl, 200 mM Tris at pH 7.4, 0.1% Tween-20) was used to block the PVDF membranes for 2 hrs. After that, they were incubated overnight at 4°C in 5% BSA/TTBS containing rabbit anti-STING (0.058 µg/ml, Cat. # 13647, RRID: AB_2732796; Cell Signaling Technology), anti-phospho-TMEM173/STING S366 (1 µg/ml, Cat. # AF7416. RRID: AB_2843856; Affinity Biosciences), anti-IFN-α (2 µg/ml, Cat. # PA5-86767, RRID: AB_2803527; Invitrogen), anti-IFN-β1 (0.48 µg/ml, Cat #. 97450, RRID: AB_2800278), anti-IRF3 (0.103 µg/ml, Cat. # 4302, RRID: AB_1904036; Cell Signaling Technology), anti-phospho-IRF3 S396 (0.046 µg/ml, Cat. # 4947, RRID: AB_823547; Cell Signaling Technology), anti-TBK1/NAK (0.19 µg/ml, Cat. #3504, RRID: AB_2255663; Cell Signaling Technology), anti-phospho-TBK1/NAK S172 (0.141 µg/ml, Cat. # 5483, RRID: AB_10693472; Cell Signaling Technology), anti-eIF4E (0.116 µg/ml, Cat. # 9742, RRID: AB_823488; Cell Signaling Technology), anti-phospho-eIF4E S209 (0.56 µg/ml Cat. # ab76256, RRID: AB_1523534; Lot: GR3263811-3, Abcam), or anti-β-Actin (0.03 µg/ml, Cat. # 4967, RRID: AB_330288; Cell Signaling Technology). Next, membranes were incubated for 2 h at room temperature in 5% BSA/TTBS containing goat-anti-rabbit IgG secondary antibody (0.04 µg/ml, Cat. # 111-035-003, RRID: AB_2313567; Lot: 158560, Jackson Immunoresearch). Protein signal detection was achieved with the chemiluminescence system (ECL plus). Bands were quantified by densitometry using an image analysis program (Image Lab Software Version 5.2.1; Bio-Rad).

### Immunohistochemistry (IHC)

Mouse DRG sections were cut from frozen blocks with a cryostat (Leica CM1950) at 20 µm thickness and mounted on to charged slides. The tissues were fixed with 10% formalin for 10 min followed by serial permeabilization with 50% ethanol, 75% ethanol and 100% ethanol for 5 min each. The slides were allowed to dry at room temperature until moisture from the slides evaporated. Hydrophobic boundaries were drawn around the sections with a Pap-pen (ImmEdge® Hydrophobic Barrier Pap-Pen, Cat. #H-4000, Vector Laboratories) and the slides were allowed to further dry for 30 min at room temperature. Sections were blocked with 10% normal goat serum with 0.3% Triton X-100 in 0.1M PB pH 7.4 for 1 h at room temperature. The sections were then incubated overnight at 4°C in blocking solution containing primary antibodies. Immunodetection of p-IRF3 and p-eIF4E was performed using rabbit anti-p-IRF3 (0.23 µg/mL, Cat. #4947, RRID: AB_823547, Cell Signaling Technology) and rabbit anti-p-eIF4E (1.12 µg/mL, Cat. #ab76256, RRID: AB_1523534, Abcam). Immunodetection of peripherin to identify neurons was performed using chicken anti-peripherin (26 µg/mL, Cat. #CPCA-Peri, RRID: AB_2284443, EnCor Biotechnology Inc.). Sections were washed in 0.1 M PB pH 7.4 and incubated for 1 h at room temperature with the corresponding secondary antibodies against p-IRF3 and p-eIF4E (0.2 µg/mL, Cat. #111-605-144, RRID: AB_2338078, Jackson ImmunoResearch Laboratories Inc.) and peripherin (0.2 µg/mL, Cat. #A11039, RRID: AB_2534096, Fisher Scientific). Sections were washed three times in 0.1 M PBS, dehydrated, and then cover-slipped with an antifade mountant (Cat. #P36390, Invitrogen). Images were captured on Olympus FV1200 Confocal Microscope System with 20X objective. Images were analyzed using Olympus CellSens Software. To determine the percentage of p-IRF3 and p-eIF4E immunoreactivity in neurons of each mouse, about 200 neurons were counted per DRG section. For the size–fluorescence data, measurements of area of peripherin, p-IRF3, and p-eIF4E in neurons were performed using a computerized image analysis system (Olympus CellSens) and plotted in 100µm^2^ (cross sectional area) increments. In addition, neuronal population with high peripherin mean fluorescence intensity (<400 µm^2^) considered as peripherin-positive, which are likely to contain nociceptors (33), were used to assess the changes in immunoreactivity of p-IRF3 and p-eIF4E. Results are reported as mean fluorescence intensity within each size population.

### FLIR Imaging

Changes in temperature either on the site of administration (tail base) or abdomen wall were assessed using a FLIR T-Series Thermal Imaging Camera (FLIR Systems, Inc). Animals were allowed to acclimate in the testing room for 1 h (ambient temperature of 21°C ± 2°C) in Plexiglas chambers (11.4 × 7.6 × 7.6 cm) before testing. We captured colorized infrared thermogram images containing the abdomen wall and tail of mice before experimental treatment and at 1, 3, 5 and 8 days after the first administration of vinorelbine. The temperature was obtained drawing a straight line at the base of the tail or a square on the abdomen of mice. The mean temperature was recorded from the average of each pixel along the drawn line or in the square. Three thermograms were averaged to obtain the mean temperature of the tail and the abdomen per animal.

### Mouse behavior

We performed behavioral experiments in wild type (WT) or transgenic male and female mice. Animals that were cohabiting were arbitrarily assigned to groups (control or treatment). All behavioral measurements were performed in age-matched animals. Animals were habituated to the experimental setup (Plexiglas chambers (11.4 × 7.6 × 7.6 cm)) for the von Frey test 1 h before each experiment. Behavioral testers were blinded to treatment and genotype in all experiments. Mechanical paw withdrawal thresholds were measured using the up-down method with calibrated von Frey filaments (Stoelting Company, Wood Dale, IL). We calculated this threshold by using the formula: 50% g threshold = 10(Xf+κδ)/10000, where Xf = the value (in log units) of the final von Frey filament used, κ = the value from look up table for the pattern of positive and negative responses published previously (34), and ∂□=□the mean difference (in log units) between stimuli. Increasing or decreasing forces with different von Frey filaments were applied during three seconds to the mouse paw in order to calculate the 50% withdrawal threshold.

Mouse grimace scoring was performed as a behavioral test for examination of spontaneous pain response (35). Mice were placed individually in a Plexiglas chamber and allowed to acclimate for 1 h, and then scored by blinded scorers at 1, 3, 5, 8, and 10 days after the first vinorelbine administration. The scores of each animal were averaged at each time point.

### Statistics

Data were analyzed with Graphpad Prism V9 (Graphpad, San Diego, CA). All data are shown as mean ± standard error of the mean (SEM). Repeated measures two-way ANOVA was used to analyze behavioral data plotted as group by time. Students t-test was used to assess effect sizes and Western blot data. Other statistical tests are described in figure legends. We used at least 5 mice per group of each sex in behavioral experiments based on power analysis predicated on our previous work with type I IFNs. Because nosex differences were observed in initial behavioral experiments with vinorelbine, we used male and female mice throughout our experiments with all data pooled from both sexes.

## Results

### Characterizing pain behavior responses induced by systemic administration of the STING activator, vinorelbine

Vinorelbine is a chemotherapeutic agent reported to induce neuropathy (10) and pain in patients (1). In our first experiments we assessed the effects of vinorelbine on pain behaviors in mice (**Fig 1A-D**). Intravenous administration of vinorelbine (day 0 and 7), but not vehicle (3% DMSO), increased the grimacing score in WT ICR male and female animals from day 1 until day 10 after administration, suggesting a sustained pronociceptive effect with a peak occurring the day after the second administration (**Fig. 1A**). Similarly, vinorelbine decreased the paw withdrawal threshold (**Fig. 1C**) during the period of evaluation, which was interpreted as mechanical nociceptive hypersensitivity. These findings can also be observed in the total effect size of grimace score and withdrawal threshold (**Fig. 1B, 1D**). Together, these experiments demonstrate that vinorelbine promotes pain-like behaviors in mice.

**Figure 1.**
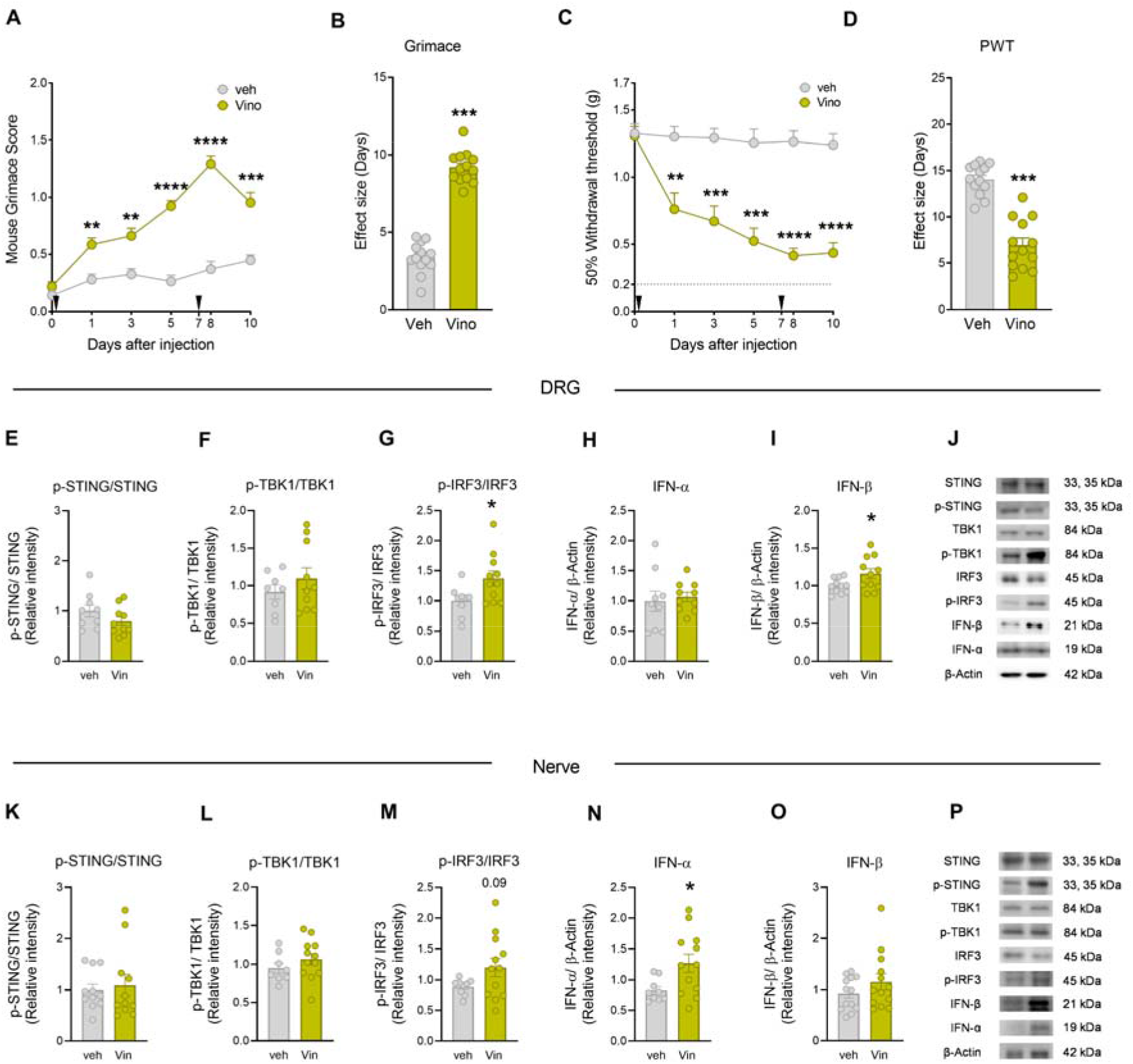
Vinorelbine induced mechanical hypersensitivity and spontaneous pain in mice associated with activation of STING-IRF3-IFN pathway in DRG and sciatic nerve. **A)** Time course of grimacing score in WT ICR mice at 1, 3, 5, 8, and 10 days after the first administration of vinorelbine (10 mg/kg, i.v.) or vehicle (3% DMSO i.v.). Arrow heads show the time of administration (days 0 and 7). **B)** Effect size (AUC) of vinorelbine-induced grimacing score in WT ICR mice. **C)** Time course of mechanical sensitivity in WT ICR mice at 1, 3, 5, 8, and 10 days after the first administration of vinorelbine (10 mg/kg i.v.) or vehicle (3% DMSO i.v.) in paw withdrawal threshold. **D)** Effect size (AUC) of vinorelbine-mediated mechanical sensitivity in WT ICR mice. **E-I)** Western blot analysis showing STING-IRF3-IFN pathway in WT DRGs from vinorelbine administered animals compared to vehicle. **J)** Representative western blot images showing STING-IRF3-IFN pathway in vinorelbine, or vehicle administered WT DRGs. **K-O)** Western blot analysis showing STING-IRF3-IFN pathway in vinorelbine treated WT sciatic nerve compared to vehicle. **P)** Representative western blot images showing STING-IRF3-IFN pathway in vinorelbine treated WT sciatic nerve compared to vehicle. Data are presented as the mean ± SEM. **p<0.01, ***p<0.001, ****p<0.0001 (n = 13 per group, with 7 male and 6 female mice) as determined by two-way ANOVA followed by Bonferroni’s test in A, C. *p<0.05, ***p<0.001 as determined by t test in B, D, E-I and K-O. Vino: vinorelbine, veh: vehicle.

Vinorelbine causes local inflammation at the injection site in humans. We measured surface body temperature with FLIR imaging as a non-invasive proxy for inflammation at the abdomen and site of injection (base of the tail). Despite there being no differences observed in temperature between treatment and vehicle groups at the abdomen (**Supp. Fig. 1A**), a significant elevation of temperature 1 and 3 days after the administration of vinorelbine at the base of the tail was observed in mice (**Supp. Fig. 1B and 1C**). This suggests that the dose of vinorelbine that we administered (29) causes pain and local inflammation in a manner that is similar to what is observed in patients (1).

### Vinorelbine-induced STING-type I interferon signaling in peripheral nerve and DRG

Type I interferon induction is predominantly controlled at the transcriptional level, wherein the main molecular mechanism controlling its production is the canonical STING-TBK1- IRF3-IFN pathway (36). We determined changes in the activation of these proteins in DRG and sciatic nerve in WT male and female mice at day 10 after the first administration of vinorelbine. We did not observe changes in the phosphorylation of STING (Ser366) (**Fig. 1E, 1K**) or the phosphorylation of TBK1 (Ser172) in WT mice at this time point (**Fig. 1F, 1L**). However, we found an increase in the phosphorylation of IRF3 (Ser396) in DRG and a trend toward increased phosphorylation in the sciatic nerve (**Fig. 1G, 1M**). We also observed an increase in the total levels of IFN-β in DRG (**Fig. 1I**), and an increase in IFN-α in sciatic nerve (**Fig. 1H**). This suggests that vinorelbine administration results in the activation of IRF3 stimulating type I IFN production in the periphery. These findings are in line with the observed vinorelbine-induced hypersensitivity in mice and with our previous work showing that type I interferons cause mechanical hypersensitivity in mice (18).

### Vinorelbine-induces p-IRF3 in DRG neurons in mice

p-IRF3 is the main downstream effector of STING inducing the transcription of *Ifna1* and *Ifnb1* genes (37, 38). We found that vinorelbine induced an increase in the phosphorylated form of IRF3 in DRG neurons compared to vehicle (**Fig. 2A**). A clear trend toward increased p-IRF3 was observed in all neuronal sizes but the increase was significant only in small and large-sized neurons (**Figure 2A-B**). Peripherin-, which is expressed only in unmyelinated cells, positive neurons had an enhancement of 3.5 times the p-IRF3 mean intensity value in DRGs from vinorelbine-treated versus vehicle mice (**Fig. 2C**). These results are consistent with our western blot experiments that showed an increase in p-IRF3 in DRG due to vinorelbine treatment and suggest that this change is mostly neuronal. Our results are in line with experiments showing that STING is localized mainly in small diameter and larger neurons, including Aβ neurons in the mouse DRG (22).

**Figure 2.**
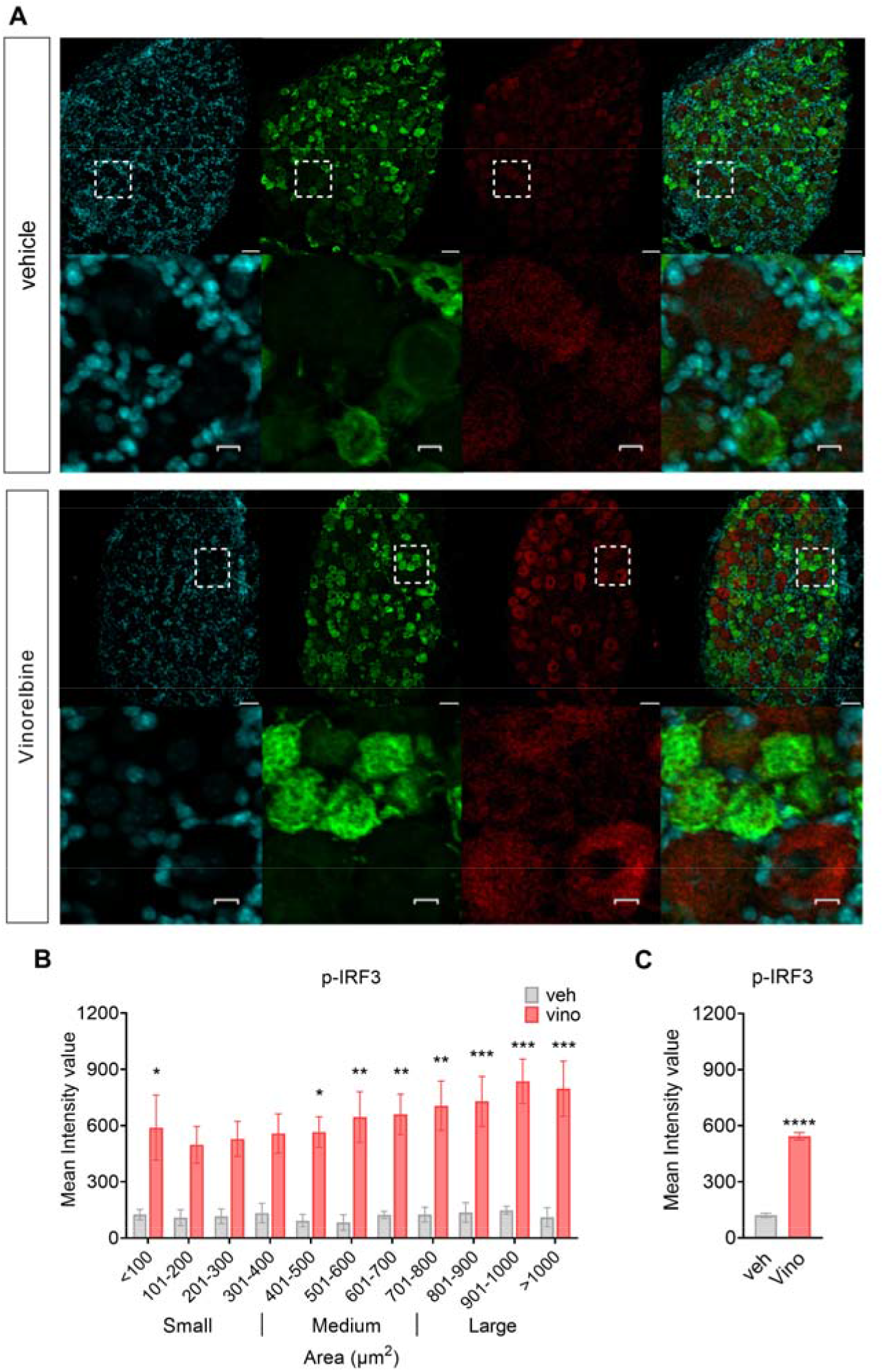
Vinorelbine increases p-IRF3 immunoreactivity in DRG neurons. **A)** Representative confocal micrographs showing p-IRF3 (red) and peripherin (green) immunofluorescence in WT ICR mouse DRG neurons day 3 post-second administration of vinorelbine (10 mg/kg i.v.) or vehicle (3% DMSO i.v.). DAPI (cyan) stains nuclei in the tissue. Respective bottom rows show zoomed-in images of a few neurons inside the white dashed-lined box. Scale bar – 10 µm. **B)** Analysis of immunoreactivity of p-IRF3 across all neuronal sizes in vinorelbine and vehicle administered WT ICR mice. The mean intensity values are plotted as a function of different neuronal sizes. **C)** Mean intensity value of p-IRF3 in peripherin-positive neurons in vinorelbine and vehicle administered WT ICR mice. Data are presented as mean ± SEM. Section thickness – 20 µm. Scale bar – 50 µm. *p<0.05, **p<0.01, ***p<0.001 (n = 3 in vehicle group, n = 4 in vinorelbine group) as determined by two-way ANOVA followed by Bonferroni’s multiple comparisons test in **B**. ****p<0.0001 as determined by t test in **C**.

### Vinorelbine-induced nociception and type I IFN induction requires STING signaling

We sought to assess whether vinorelbine-induced nociceptive effects depend on STING activation. To this end, we took advantage of the mutant mouse strain, Goldenticket (Sting*^Gt/Gt^*) also known as *Tmem173^Gt^*, that fails to produce type I IFNs upon activation with c-di-GMP or infection with pathogens (39, 40). Sting*^Gt/Gt^* animals harbor a chemically induced (N-ethyl-N-nitrosourea) mutation in exon 6 of *Sting* resulting in an isoleucine to asparagine change in amino acid 199 located in the carboxyl-terminal end (CTT) of the protein, which is an important site to recruit TBK1, the kinase that phosphorylates IRF3, thus signaling via IRF3 is lost in Sting*^Gt/Gt^* mice. As expected, western blotting experiments revealed a profound reduction (78.9% in DRG and 76.9% in sciatic nerve) in the levels of STING in Sting*^Gt/Gt^* mice, which render STING barely detectable (**Supp. 2A-B**).

We assessed the effects of vinorelbine in Sting*^Gt/Gt^* mice and their WT controls. Because no sex differences were observed in previous experiments, male and female mice were used as part of the same cohort in these experiments. As expected, vinorelbine induced an increase in grimace score in WT mice from day 5 after first administration (day 0) and lasting until day 10. However, vinorelbine had no effect in Sting*^Gt/Gt^* mice (**Fig. 3A**). This genotype difference was also observed in the effect size (**Fig. 3B**). Additionally, a reduction in the paw withdrawal threshold was observed in WT mice that was not observed with vinorelbine in Sting*^Gt/Gt^* mice (**Fig. 3C-D**). These results indicate that STING activation is needed for the pronociceptive effects of vinorelbine in mice.

**Figure 3.**
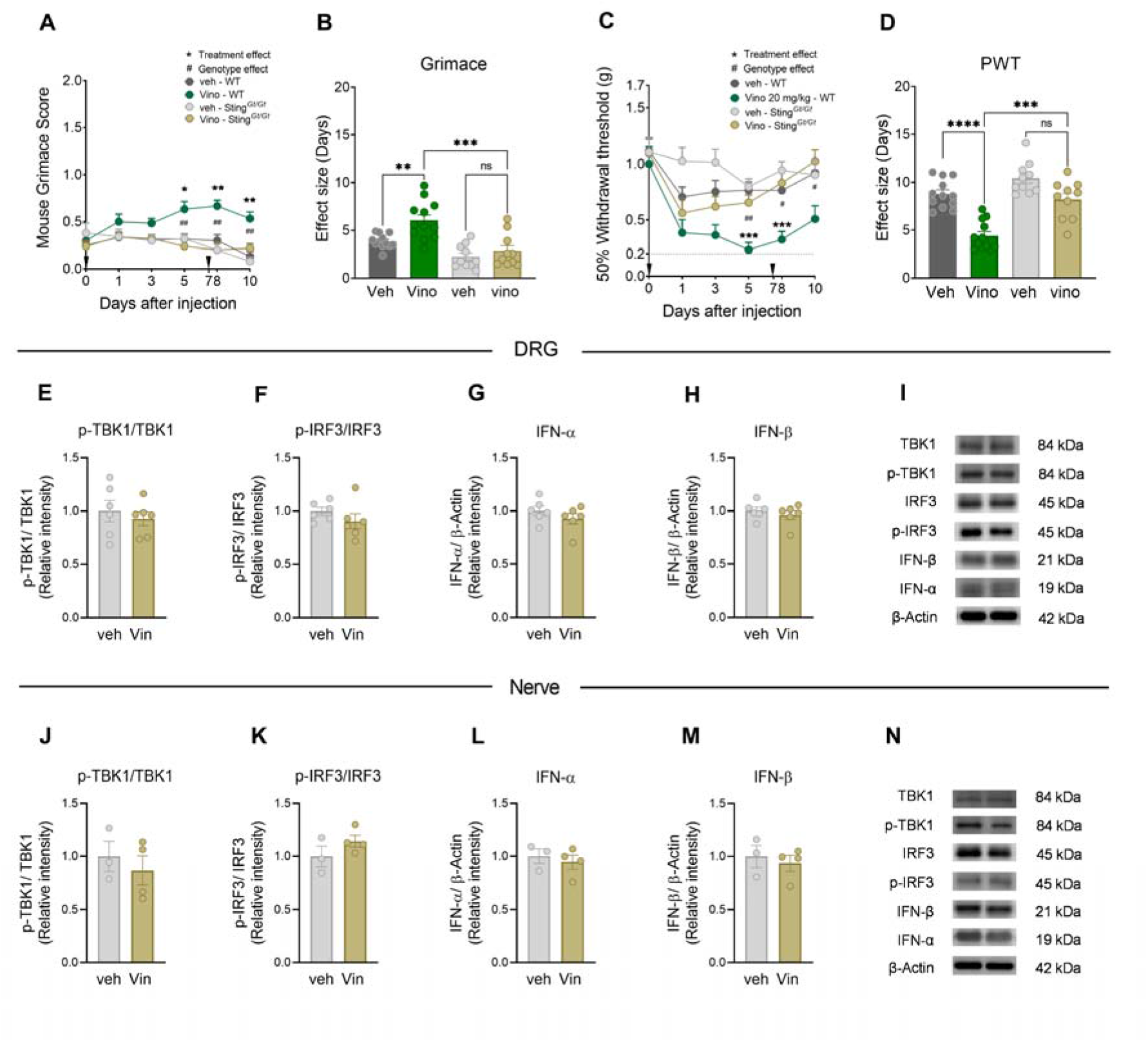
Sting*^Gt/Gt^* mice show decreased vinorelbine induced nociception, and lose p-IRF3 and type I IFN induction in peripheral nerves. **A)** Time course of grimacing score in Sting*^Gt/Gt^* and C57BL/6J WT mice at 1, 3, 5, 8, and 10 days after the first administration of vinorelbine (10 mg/kg, i.v.) or vehicle (3% DMSO i.v.). Arrow heads show the time of administration (days 0 and 7). **B)** Effect size (AUC) of vinorelbine-induced grimacing score in Sting*^Gt/Gt^* and C57BL/6J WT mice. **C)** Time course of mechanical threshold in Sting*^Gt/Gt^*and WT C57BL/6J mice at 1, 3, 5, 8, and 10 days after the first administration of vinorelbine (10 mg/kg i.v.) or vehicle (3% DMSO i.v.) in paw withdrawal threshold. **D)** Effect size (AUC) of vinorelbine-mediated mechanical sensitization on paw withdrawal threshold in Sting*^Gt/Gt^* and C57BL/6J mice. **E-I)** Western blot analysis showing STING downstream pathway in DRGs from vinorelbine administered Sting*^Gt/Gt^* mice compared to vehicle. **J)** Representative western blot images showing STING downstream pathway in DRGs from vinorelbine administered Sting*^Gt/Gt^*mice compared to vehicle. **K-O)** Western blot analysis showing STING downstream pathway in sciatic nerve from vinorelbine administered Sting*^Gt/Gt^* mice compared to vehicle. **P)** Representative western blot images showing STING downstream pathway in sciatic nerve from vinorelbine administered Sting*^Gt/Gt^* mice compared to vehicle. Data are presented as the mean ± SEM. *p<0.05, **p<0.01, ***p<0.001 vs Vino Sting*^Gt/Gt^*, and ^#^p<0.05, ^##^p<0.01 vs Vino C57BL/6J (n = 10-12 per group, 6 males and 4-6 females per group) as determined by two-way ANOVA followed by the Bonferroni’s test in **A,C**. ***p<0.001, ****p<0.0001 as determined by two-way ANOVA followed by Bonferroni’s test in **B,D**. t test in **E-I** and **K-O** (n=3-6 mice per group). Vino: vinorelbine, veh: vehicle.

We performed Western blots to further elucidate the changes of the canonical STING-IFN pathway underlying the blunted hypersensitivity of vinorelbine in Sting*^Gt/Gt^*mice. In the absence of a functional STING pathway, the administration of vinorelbine was not able to result in the increased production of IFN or p-IRF3 in DRG and sciatic nerve (**Fig. 3E-P**), which aligns with the lack of hypersensitivity induced by the chemotherapeutic in Sting*^Gt/Gt^* mice. These results support the conclusion that vinorelbine induces type I IFN production via STING activation, consistent with the loss of nociceptive sensitization in Sting*^Gt/Gt^* mice.

### Loss of type I IFN-MNK-eIF4E signaling reduces the vinorelbine-elicited pain in MNK1 KO mice

MNK1 is a serine/threonine kinase that phosphorylates eIF4E, the 5’ mRNA Cap-binding protein, and is a key contributing factor in the development of chronic pain (41-44). Previously, we showed that type I IFNs stimulate ERK/MAP kinase-MNK-eIF4E signaling in mouse DRG neurons (18). Since vinorelbine promotes the production of type I IFNs in DRG and sciatic nerve, we tested the hypothesis that vinorelbine-evoked nociceptive behaviors require MNK1 signaling. Vinorelbine was administered to MNK1 KO mice and their wild type controls (C57BL/6J) and compared its effects with vehicle in behavioral assays. As expected, we found an increase in the grimace score in WT mice from day 1 after administration to day 10 (**Fig. 4A**). A reduction in the vinorelbine-induced pronociceptive effect was observed in MNK1 KO mice with a significantly lower grimace score the day after the second administration compared to WT animals (**Fig. 4A**). The statistical difference in the genotype effect can also be observed in the effect sizes in vinorelbine treated animals (**Fig. 4B**). Similarly, we found a reduction of paw withdrawal threshold in WT animals that reached almost 0.2 g at its peak. While a significant decrease in von Frey threshold was observed in the MNK1 KO mice 1 day after treatment, this effect was short lived and not seen on subsequent days (**Fig. 4C**). In line with this, the paw withdrawal threshold effect sizes of vinorelbine were significantly greater in WT animals compared to MNK1 KO mice (**Fig. 4D**).

**Figure 4.**
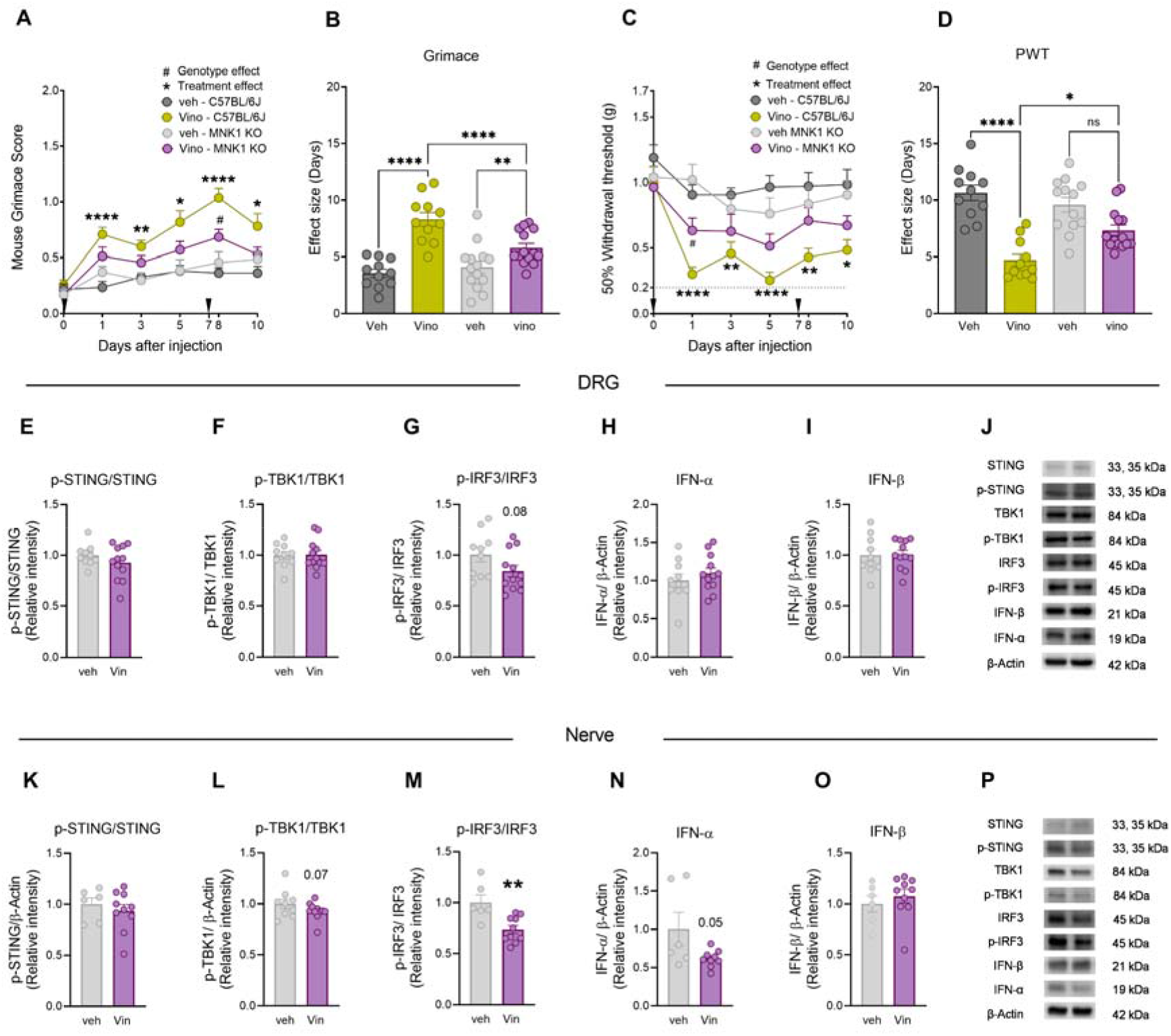
The lack of MNK1 impairs vinorelbine-induced pain and STING-IRF3-IFN activation pathway. **A)** Time course of grimacing score in *MNK1 KO* and C57BL/6J WT mice at 1, 3, 5, 8, and 10 days after the first administration of vinorelbine (10 mg/kg, i.v.) or vehicle (3% DMSO i.v.). Arrow heads show the time of administration (days 0 and 7). **B)** Effect size (AUC) of vinorelbine-induced grimacing score in *MNK1 KO* and C57BL/6J WT mice. **C)** Time course of allodynia in *MNK1 KO* and WT C57BL/6J mice at 1, 3, 5, 8, and 10 days after the first administration of vinorelbine (10 mg/kg i.v.) or vehicle (3% DMSO i.v.) in paw withdrawal threshold. Arrow heads show the time of administration (days 0 and 7). **D)** Effect size (AUC) of vinorelbine-mediated allodynia on paw withdrawal threshold in *MNK1 KO* and C57BL/6J mice. **E-I)** Western blot analysis showing STING-IRF3-IFN pathway changes in DRGs from vinorelbine administered *MNK1 KO* mice compared to vehicle. **J)** Representative western blot images showing STING-IRF3-IFN pathway in DRGs from vinorelbine administered *MNK1 KO* mice compared to vehicle. **K-O)** Western blot analysis showing STING-IRF3-IFN pathway in sciatic nerve from vinorelbine administered *MNK1 KO* mice compared to vehicle. **P)** Representative western blot images showing STING-IRF3-IFN pathway in DRGs from vinorelbine administered *MNK1 KO* mice compared to vehicle. Data are presented as the mean ± SEM. *p<0.05, **p<0.01, ****p<0.0001 vs Vino *MNK1 KO*, and #p<0.05 vs Vino C57BL/6J (n = 10-14 per group, 5-7 mice per sex per group) as determined by two-way ANOVA followed by the Bonferroni’s test in **A,C**. *p<0.05, **p<0.01, ****p<0.0001, as determined by two-way ANOVA followed by Bonferroni’s test in **B,D**. **p<0.01 as determined by t test in **E-I** and **K-O**. Vino: vinorelbine, veh: vehicle.

Considering the reduction of the pronociceptive effects of vinorelbine in MNK1KO, in an independent set of experiments we measured the effects of vinorelbine on STING-TBK1- IRF3-IFN signaling. Vinorelbine treatment was associated with a trend toward decreased levels of p-TBK1 and p-IRF3 in MNK1 KO mice (**Fig. 4F-G**) in DRG, and it did not modify the levels of p-STING (**Fig. 4E**) or type I IFN production (**Fig. 4H-I**). Vinorelbine provoked similar changes in the STING-TBK1-IRF3 pathway in sciatic nerve decreasing p-TBK1 and p-IRF3 (**Fig. 4M**), but in addition the drug also downregulated IFN-α levels in MNK1 KO mice (**Fig. 4N**). Together, these results show an interplay between the STING-IRF3-IFN pathway and MNK1 signaling, highlighting that vinorelbine fails to induce increases in IFN production in peripheral nerves in the absence of MNK1.

Because we observed a reduction in the pro-nociceptive effect of vinorelbine in MNK1 KO animals as well as abrogated associated changes in the STING pathway, we investigated the effect of vinorelbine on p-eIF4E in WT, Sting*^Gt/Gt^*, and MNK1 KO animals. Moreover, translation regulation through p-eIF4E has been shown to be associated with antiviral response and translation of mRNAs encoding proteins such as type I IFNs (45). An increase in p-eIF4E in DRG neurons was observed across all neuronal sizes (**Fig. 5A-B**). Specifically, peripherin-positive neurons from vinorelbine administered animals had an increase in p-eIF4E of 0.4 times the mean intensity value compared to control (**Fig. 5C**). To give more insight about p-eIF4E changes induced by vinorelbine, western blot experiments in animals of the different genotypes were performed in DRG and sciatic nerve. In line with our immunohistochemistry experiments we found a clear trend toward an increase of p-eIF4E in DRG from wild type animals (**Fig. 6A**). On the other hand, no changes were found in DRGs of Sting*^Gt/Gt^* and MNK1 KO mice (**Fig. 6B-C**). An increase in p-eIF4E was observed in sciatic nerve in wild type mice (**Fig. 6D**). Administration of vinorelbine resulted in no change in p-eIF4E levels in Sting*^Gt/Gt^* in sciatic nerve (**Fig. 6E**). Further, a trend toward reduction in p-eIF4E with vinorelbine administration was observed in MNK1 KO sciatic nerve (**Fig. 6F**). Both MNK1 and MNK2 are expressed in mouse DRG neurons (46) so the remaining p-eIF4E in peripheral nerves in MNK1 KO mice is likely support by the MNK2 isoform, which is known to have constitutive activity (26).

**Figure 5.**
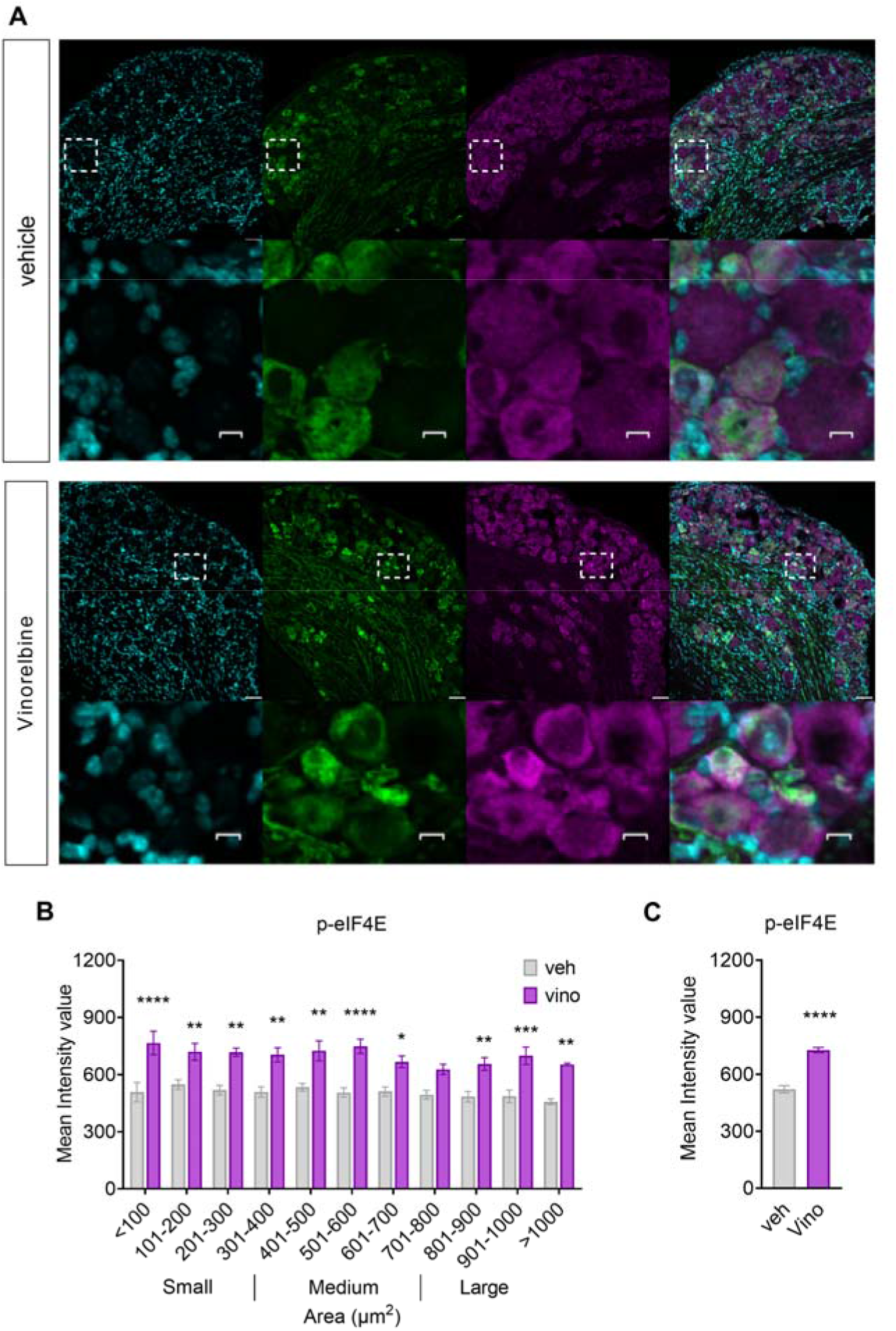
Vinorelbine increases p-eIF4E in WT DRG neurons and sciatic nerve but not in Sting*^Gt/Gt^*, and MNK1 KO mice. **A)** Representative confocal micrographs showing p-eIF4E (magenta) and peripherin (green) immunofluorescence in WT mouse DRG neurons at day 3 post-second administration of vinorelbine (10 mg/kg, i.v.) or vehicle (3% DMSO i.v.). DAPI (cyan) stains nuclei in the tissue. **B)** Analysis of immunoreactivity of p-eIF4E across all neuronal sizes in vinorelbine and vehicle administered WT mice. The mean intensity values are plotted as a function of different neuronal sizes. **C)** Mean intensity value of p-eIF4E in peripherin-positive neurons in vinorelbine and vehicle administered WT mice. Data are presented as mean ± SEM. Section thickness – 20 µm. Scale bar – 50 µm. Respective bottom rows show zoomed in images of a few neurons inside the white dashed-lined box. Scale bar – 10 µm. *p<0.05, **p<0.01, ***p<0.001, ****p<0.0001 (n = 3 per group) as determined by two-way ANOVA followed by Bonferroni’s multiple comparisons test in **B**. Data are presented as mean ± SEM ****p<0.0001 as determined by t test in **C**.

**Figure 6.**
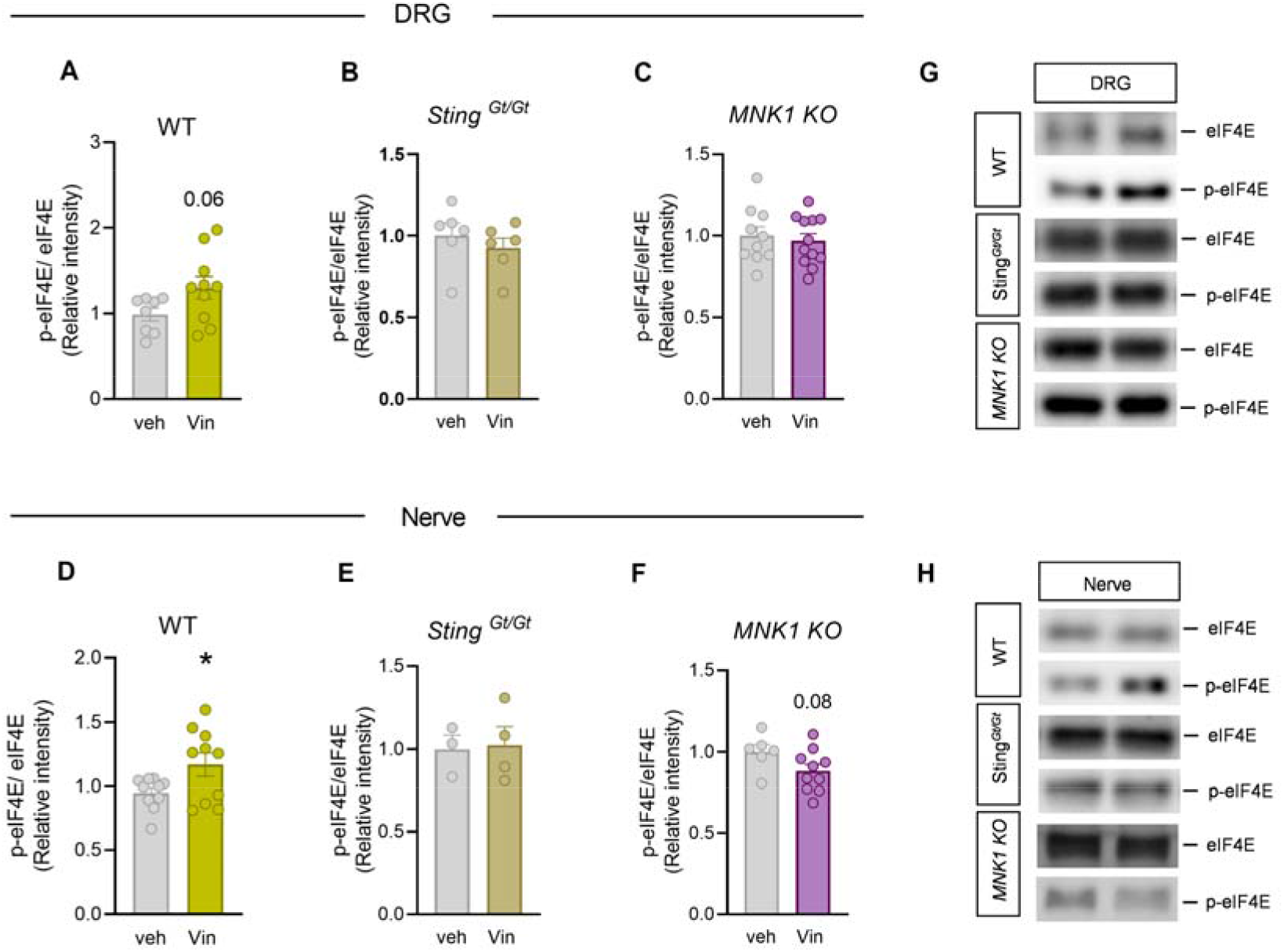
Vinorelbine enhances p-eIF4E in WT DRG and sciatic nerve but not in Sting*^Gt/Gt^*, and MNK1 KO mice. **A-C)** Western blot analysis of p-eIF4E in DRGs of WT, Sting*^Gt/Gt^*, and *MNK1 KO* vinorelbine-or vehicle-administered mice. **D-F)** Western blot analysis of p-eIF4E in sciatic nerve of WT, Sting*^Gt/Gt^*, and *MNK1 KO* vinorelbine-or vehicle-administered mice. **G)** Representative western blot images showing p-eIF4E bands in DRGs of WT, Sting*^Gt/Gt^*, and *MNK1 KO* vinorelbine-or vehicle-administered mice. **H)** Representative western blot images showing p-eIF4E mean intensity levels in sciatic nerve of WT, Sting*^Gt/Gt^*, and *MNK1 KO* vinorelbine-or vehicle-administered mice. Data are presented as mean ± SEM *p<0.05 as determined by t test in **D**.

Altogether, our results reinforce the idea that IFNs cause pain by acting on nociceptors and explain, mechanistically, how chemotherapeutics that are STING activators and microtubule destabilizers such as vinorelbine, have the potential to cause pain and peripheral neuropathy. Furthermore, this study gives insight into the complexity of IFN induction within peripheral nerves and provides additional evidence that translational regulatory mechanisms play a critical role in chemotherapy induced pain (41).

## Discussion

We reach several primary conclusions from the experiments described here. First, we have discovered a plausible mechanism through which vinorelbine induces pain hypersensitivity in male and female mice by activating the STING signaling pathway, including IRF3 phosphorylation (S396), and subsequent type I IFN induction in the peripheral nervous system. This conclusion is supported by the finding that vinorelbine-induced pain is abrogated in Sting*^Gt/Gt^* mice and peripheral nerve induction of type I IFN by vinorelbine was absent in these mice. Second, our work shows that a downstream effector involved in vinorelbine-induced pain is MNK1-eIF4E signaling, a finding consistent with our previous work on mechanisms through which type I IFNs cause pain in mice (18). Therefore, our work also provides a clear mechanistic understanding of vinorelbine-induced peripheral neuropathy and leads us to the third conclusion, that MNK signaling is a therapeutic avenue for reducing the neuropathy caused by this chemotherapeutic. Future work will be aimed at understanding how MNK inhibition can be used as a therapeutic approach to reduce chemotherapy neuropathies and improve cancer chemotherapeutic efficacy.

Our work was primarily motivated by the consistent clinical finding that vinorelbine causes painful peripheral neuropathy in humans treated with this drug. Our work fills an important gap in knowledge by providing a mechanistic explanation for vinorelbine-induced pain. Neuropathy has been observed in oncological patients receiving vinorelbine in phase II/III trials for metastatic breast cancer, with neurotoxicity ranging from 33 to 43.6% (47, 48). Notably, a heightened severity of neurotoxicity (87%) after 4 cycles of vinorelbine has also been reported (6). Patients who undergo treatment with vinorelbine report pain, primarily localized at the tumor site or in regions affected by nerve compression caused by the tumor (1, 47, 49). Moreover, some patients also present with paresthesia and myalgia consisting of acute pain in lower back muscles and extremities (47). There is also evidence that vinorelbine causes a peripheral neuropathy because the drug decreases the amplitude of evoked action potentials in sensory nerve bundles in patients (6), one of the main neurophysiological abnormalities observed in neuropathy (50). Finally, vinorelbine treatment is associated with the worsening of peripheral neuropathy in patients with pre-existing diabetes or CIPN (51). Despite the induction of neuropathy, vinorelbine is approved for advanced NSCLC treatment (4) and is used in clinical trials for a wide variety of cancer types (1, 5, 47, 48). Based on the rationale that vinorelbine causes neuropathic pain and is used alone or in combination with other chemotherapeutic drugs (52), we studied the signaling pathways associated with vinorelbine-related pain to find new strategies to mitigate or prevent the concomitant pain associated with its use. Our results provide a detailed mechanistic rationale for blocking MNK signaling as a therapeutic strategy to mitigate the pain-promoting effects of vinorelbine.

A second motivation for our work was to better understand the role of STING and type I IFN signaling in painful peripheral neuropathy. Recently, it was demonstrated that microtubule-targeted agents (MTAs), including vinorelbine and eribulin, activate the STING pathway leading to type I IFN production in human breast cancer and leukemia monocytic cell lines (12, 28). The STING pathway is an intracellular sensor system of external or self-origin DNA that is critical for recognizing pathogen infection and/or cellular damage to establish an effective host defense (53, 54). Our results clearly demonstrate that vinorelbine induces STING activation in mouse DRG neurons and peripheral nerve as reflected in increased IRF3 phosphorylation and type I IFN induction (55). Recently published work shows that STING induces peripheral nociceptive sensitization via NFKB activation in DRG neurons in a model of bone cancer pain in rats (25). Furthermore, the STING inhibitor C-176 has been shown to reduce mechanical hypersensitivity induced by nerve injury in rats suggesting a pronociceptive role for STING (24). Mounting evidence has shown that STING is anomalously activated in models of peripheral neuropathic pain (56), central neuropathic pain (57, 58), and inflammatory pain (25, 59) in rodents. These works have concluded that the pharmacological blockade of STING may be a promising target for pain (25, 56-59). Collectively, our work is consistent with the emerging view that activation of STING signaling in the peripheral nervous system is pro-nociceptive and may play an important role in multiple types of painful peripheral neuropathies.

Evidence of an antinociceptive role of STING has also been reported (22, 23, 60). Activators of STING, including ADU-S100, DMXAA and 3’3’-cGAMP, have been shown to reverse CIPN-, nerve injury- (22) and bone cancer-induced pain in mice (23) via suppression of nociceptor excitability. Under physiological conditions, the role of STING has been shown to regulate steady-state nociception, since intrathecal administration of the STING agonists ADU-S100 or DMXAA produced antinociception in naïve mice. Moreover, naïve Sting*^Gt/Gt^*mice showed increased sensitivity to mechanical and cold stimuli, although we did not find this in our experiments for von Frey threshold, and DRG nociceptors from these mice displayed increased action potential firing and reduced rheobase suggesting nociceptor hyperexcitability (22). Thus, the aforementioned studies suggest an antinociceptive role for STING (23). A recent review concluded that STING signaling causes bidirectional effects on nociception but did not reach firm conclusions on why STING activation may be associated with increased pain in some contexts and anti-nociception in others (54). One hypothesis to explain this controversy is that whether STING is activated in the peripheral or central nervous system (CNS) is critical for the direction of the effects of the pathway on nociceptive signaling. From this perspective, it is important to note that the chemical properties of vinorelbine, like other vinca alkaloids such as vincristine, render the drug unable to cross the blood-brain barrier (61). Accordingly, we only observed pro-nociceptive effects of vinorelbine in mice.

There is controversy over the effects of type I IFNs on nociception, which may also be explained by peripheral nervous system (PNS) versus CNS sites of action (62). We have previously shown that type I IFNs activate their receptors (IFNR) localized on nociceptors driving neuron hyperexcitability and pro-nociceptive actions in mice (18). Likewise, systemic IFN-α increases formalin-evoked nociceptive behavior in mice (20). In the same line, patients with hepatitis undergoing IFN-α therapy reported experiencing pain with IFN infusion (21). On the other hand, other studies have shown that STING-type I IFN signaling attenuates DRG neuron excitability via suppression of Na^+^ and Ca^2+^ channels activity (22, 23), suggesting an antinociceptive role for IFNs. Notably, IFN-α administered in the periphery (i.pl.) led to mechanical hypersensitivity, which was alleviated by intrathecal administration of IFN-α (22), again consistent with different actions of this signaling pathway depending on the site of administration. Other studies finding an antinociceptive role for type I IFNs have administered the IFNs intracerebroventricularly or intrathecally (63-65). Our work adds to the growing body of evidence that peripheral type I IFN signaling is pro-nociceptive. We did not test CNS-mediated effects of vinorelbine because this is not relevant for the clinical effects of this peripherally-restricted drug.

Findings in Sting*^Gt/Gt^* mice unequivocally implicate STING signaling in the pro-nociceptive effects of vinorelbine. Accordingly, we did not observe changes in STING downstream effectors in either the peripheral nerve or DRG with vinorelbine administration. In contrast, in WT mice, we consistently observed increased IRF3 phosphorylation using Western blotting and immunofluorescence. We also observed increases in type I IFNs in DRG and sciatic nerve in WT mice, which likely occurred due to the presence of IRF3-driven transcription of *Ifna* and *Ifnb* genes (37, 38). On the other hand, we did not find changes in the phosphorylation of STING (Ser366), which is essential to trigger STING pathway signaling (66). A possible explanation is that after trafficking to the Golgi apparatus, STING is phosphorylated by serine/threonine UNC-51-like kinase (ULK1/ATG1) to facilitate its degradation and prevent the deleterious effect of a long-lasting inflammatory state induced by persistent STING activation (67). Thus, our failure to observe changes in phosphorylation of STING could be due to degradation of the activated protein at the timepoint at which we harvested tissues for analysis. This could also explain the lack of changes in p-TBK1 in WT mice.

The exact mechanism by which the MTAs such as vinorelbine or eribulin enhance STING signaling has not been fully uncovered. Some studies have suggested that the STING pathway is activated by mitochondrial or nuclear DNA release induced by cellular damage resulting from the treatment with MTAs but it has also been proposed that STING activation occurs downstream of the microtubule disruption triggered by these chemotherapeutics (12, 28, 68). Whatever the precise mechanism turns out to be, the discovery that vinorelbine-induced pain depends on STING-type I IFN signaling makes it possible to implement strategies to alleviate the neuropathic pain state caused by the chemotherapeutic. We demonstrate that this can be achieved by targeting MNK1-eIF4E signaling. Vinorelbine caused an increase in eIF4E phosphorylation at a site that is specifically regulated by MNK in both the peripheral nerve and in DRG cell bodies. Moreover, grimacing and mechanical hypersensitivity were profoundly reduced in MNK1 KO mice. Translation regulation via MNK-eIF4E signaling has been shown to participate in the development and maintenance of pain hypersensitivity in many different preclinical models (18, 41, 42, 44). MNK1 has been shown to be expressed at the mRNA level in nearly all human nociceptors (46) indicating that this mechanism is likely to translate to humans. Our findings support the idea that MNK represents a promising mechanistic target for pharmacological relief or prevention of chemotherapy-induced pain. It will be important to understand whether MNK inhibition will interfere with the anti-cancer activity of the chemotherapeutic, but this is unlikely since increased MNK activity and increased eIF4E phosphorylation is associated with many cancers (69, 70) and MNK inhibitors are currently in clinical trials for cancer treatment (71).

There are several limitations of this study. We did not directly interfere with type I IFNs, which would have allowed us to dissect STING activation versus type I-IFN downstream effects. While this can be addressed in future studies, our main goal was to understand whether STING activation was linked to vinorelbine-induced pain and our use of Sting*^Gt/Gt^* mice provides a clear link between vinorelbine and STING signaling. In some cases, we found trends for signaling changes in sciatic nerve or DRG (*p* values between 0.06 and 0.09). These trends were consistent with findings using transgenic and knockout mice for signaling pathways and were observed in a complex tissue where there are a mix of cell types that contribute to the overall changes in protein levels. We do not think that these observed trends weaken the overall conclusions of the study. Unanswered questions remain to be uncovered regarding other STING activated pathways, which we did not examine. For instance, it is known that STING signaling can activate inhibitor of κB kinase (IKKε)/nuclear factor κB (NF-κB), which synergizes with IRF3 to induce higher levels of type I IFNs (72). Future studies can address nuances of these signaling effects now that our work has unequivocally demonstrated that vinorelbine-induced pain requires STING signaling. Finally, while we provide genetic evidence for MNK1-eIF4E signaling as a causative factor in vinorelbine-evoked pain, it will be interesting to assess the extent to which MNK inhibitors like eFT508 can reproduce this effect. Given that pharmacological inhibition of MNK has thus-far matched genetic manipulation in behavioral endpoints in neuropathic and other pain models (18, 41, 42, 44, 73, 74) it is likely that these MNK inhibitors will be effective in reducing pain promoting effects of vinorelbine.

## Acknowledgements

We acknowledge Zawge Johannes Daniel and Abdulfetah Abdo for the technical support to this project. This work was supported by NIH grant NS065926 to TJP.

