## Supplemental Information for "Vinorelbine causes a neuropathic pain-like state in mice via STING and MNK1 signaling associated with type I interferon induction"

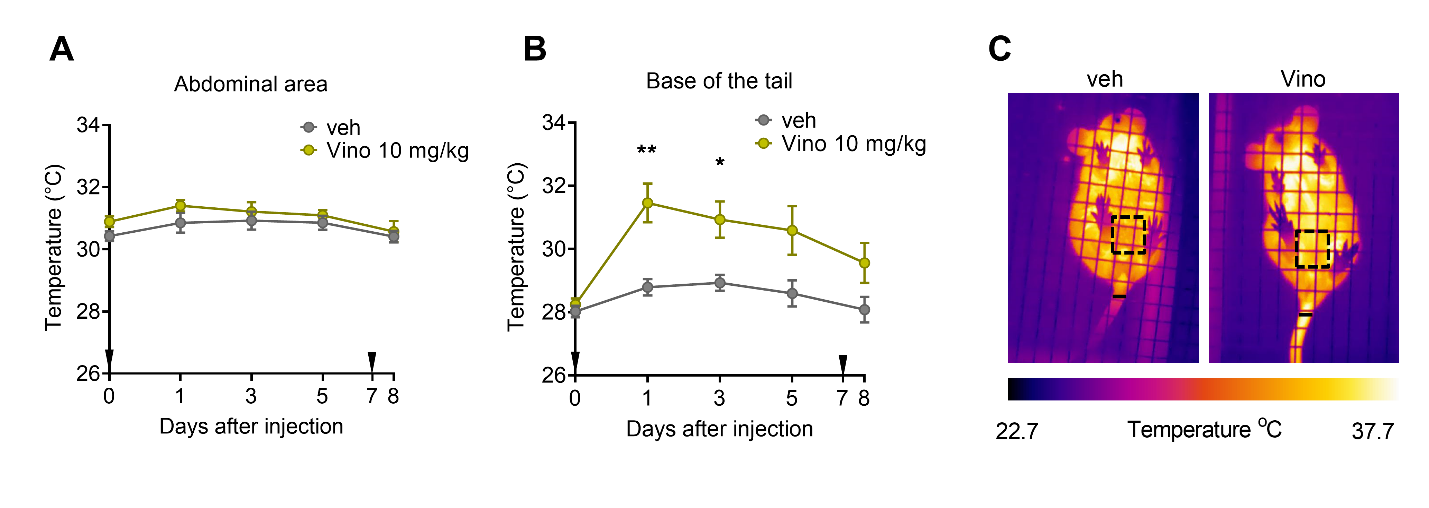

**Supplementary Fig. 1.** Changes in temperature either on the abdominal area (**A**) or the site of administration (base of tail, **B**) were assessed before experimental treatment and at 1, 3, 5 and 8 days after the first administration of vinorelbine. Representative colorized infrared thermogram images containing the abdomen wall and tail of mice on day 1 are shown in panel **C**. The color bar represents the temperature in °C. The dotted square represents the abdominal evaluated area, and the straight line represents the selected area considered at the base of the tail. Data are presented as the mean ± SEM. *p<0.05, **p<0.01 (n = 11 per group with 5 male and 6 female mice) as determined by two-way ANOVA followed by Bonferroni’s test in **A, B.** Vino: vinorelbine, veh: vehicle.

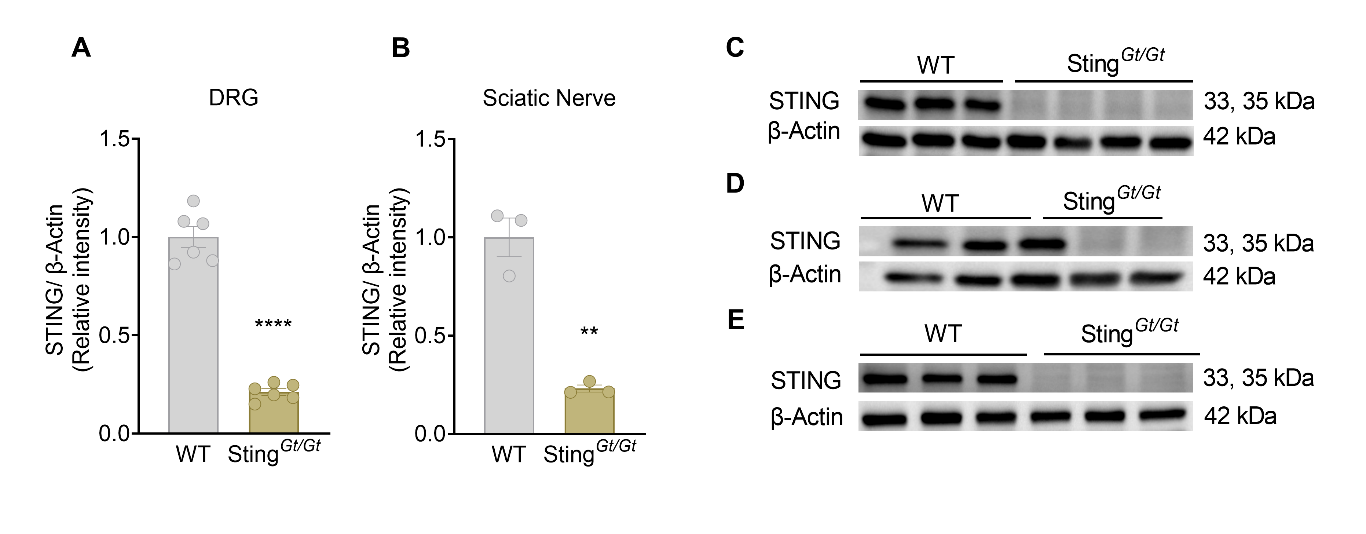

**Supplementary Fig. 2.** Sting*^Gt/Gt^* mice have a robust reduction in STING compared to WT mice in DRGs (**A**) as well as sciatic nerve (**B**). Representative western blot images showing STING mean intensity levels in DRGs in Sting*^Gt/Gt^* compared to WT male (**C**) and female (**D**) mice. Representative western blot images showing STING mean intensity levels in sciatic nerve in Sting*^Gt/Gt^* compared to WT mice (**E**). Data are presented as mean ± SEM. **p<0.01, ****p<0.0001 (n=6 per group in **A**, n=3 per group in **B**), as determined by unpaired t test.
